# Metagenomic diagnosis and pathogenic network profile of SARS-CoV-2 in patients co-morbidly affected by type 2 diabetes

**DOI:** 10.1101/2021.02.23.432535

**Authors:** Hassan M. Al-Emran, M. Shaminur Rahman, Md. Shazid Hasan, A. S. M. Rubayet Ul Alam, Ovinu Kibria Islam, Ajwad Anwar, Iqbal Kabir Jahid, M. Anwar Hossain

## Abstract

**Background:** The mortality of COVID-19 disease is very high among males or elderly or individuals having comorbidities with obesity, cardiovascular diseases, lung infections, hypertension, and/or diabetes. Our study characterizes SARS-CoV-2 infected patients’ metagenomic features with or without type 2 diabetes to identify the microbial interactions associated with its fatal consequences.

**Method:** This study compared the baseline nasopharyngeal microbiome of SARS-CoV-2 infected diabetic and non-diabetic patients with controls adjusted with age and gender. The mNGS were performed using Ion GeneStudio S5 Series and the data were analyzed by the Vegan-package in R.

**Results:** All three groups possessed significant bacterial diversity and dissimilarity indexes (p<0.05). Spearman’s correlation coefficient network analysis illustrated 183 significant positive correlations and 13 negative correlations of pathogenic bacteria (r=0.6-1.0, p<0.05), and 109 positive correlations among normal-flora and probiotic bacteria (r>0.6, p<0.05). The SARS-CoV-2 diabetic group exhibited a significant increase of pathogens (p<0.05) and opportunistic pathogens (p<0.05) with a simultaneous decrease of normal-flora (p<0.05). The molecular docking analysis of Salivaricin, KLD4 (alpha), and enterocin produced by several enriched probiotic strains presented strong binding affinity with Shiga toxin, outer membrane proteins (ompA, omp33) or hemolysin.

**Conclusion:** The dysbiosis of the bacterial community might be linked with severe consequences of COVID-19 infected diabetic patients, although few probiotic strains inhibited numerous pathogens in the same pathological niches. This study suggested that the promotion of normal-flora and probiotics through dietary changes and reduction of excessive pro-inflammatory states by preventing pathogenic environment might lead to a better outcome for those co-morbid patients.

## Introduction

Severe acute respiratory syndrome coronavirus-2 (SARS-CoV-2) is the etiological root of the COVID-19 pandemic, which has affected over 83 million people worldwide in 2020 ^1^. The virus expresses itself with highly variable severity, ranging from no outward symptoms to severe respiratory distress ^2^. One of the most common questions in the scientific community, why some patients are asymptomatic and others (especially co-morbid patient) have fatal consequences, remains largely unknown. The mortality rate of COVID-19 is very high among males or elderly or individuals having comorbidity with obesity, cardiovascular diseases, lung infections, hypertension, and/or diabetes. The SARS-CoV-2 mortality was 50% in males and 43% in females among ICU patients in Lombardy, Italy ^3^. The incidence of mortality is 12.2/1000 among male ICU patients than 9.9/1000 among female ICU patients per day in that city. SARS-CoV-2 infected diabetic patients’ mortality is 7.8% compared to 2.7% in non-comorbid patients in China ^4^. In Italy, 29.8% of the SARS-CoV-2 infected diabetic patients died in May 2020. The study found that the risk factors of death increased by 3.2 times from SARS-CoV-2 infection when associated with diabetes ^5^. Considering the transmission rate of this infection and that over 463 million people are already afflicted with hyperglycemia ^6^, adequate research into management of SARS-CoV-2 infection in diabetic patients is required for public safety. Almost 2 million people died in 2020 so far, and the outcome of this pandemic can be far more disastrous than that. The genetic sequences of SARS-CoV-2 is largely analyzed. As of 31^st^ of December 2020, a total of 254,153 whole-genome sequences of SARS-CoV-2 have been performed ^7^; however, the pathogenicity, carriage information, and the interactions with commensals or opportunistic bacteria of this virus are still unclear. What’s more, the development of respiratory coinfections is reported to be associated with COVID-19 disease severity and fatality in many cases ^8, 9, 10^.

Metagenomic analysis is popularly used to understand the diversity and pathogenesis of microbial populations in a group of subjects. However, metagenomics based on next-generation sequencing (mNGS) is rarely applied to the clinical samples of SARS-CoV-2 ^4, 11, 12^. This technology can be used to uncover all microbiome interacting in the same pathological niche which leads to ultimate pathophysiological conditions. The present study aims to perform meta-transcriptomic and in-depth bioanalysis to characterize SARS-CoV-2 diabetic and non-diabetic patients compared with controls to identify the microbial interaction associated with the fatal consequences.

## Methods

### Patients

Seven SARS-CoV-2 positive samples and four SARS-CoV-2 negative controls were included in this study. All positive cases were selected from the continuous surveillance at the Genome Center, Jashore University of Science and Technology covering four districts of Bangladesh, Jashore, Jhenaidah, Magura, and Narail authorized by the Directorate General of Health Services, Bangladesh, for the screening of COVID-19. Among those seven positive cases, three patients had a history of type 2 diabetes with two reported deaths. Four age and sex-matched subjects were included in this study, as healthy (N=2) controls and unknown etiology controls (N=2), confirmed as SARS-CoV-2 negative by rt-PCR. Both healthy controls have no history of fever or other illness or uptake of antibiotics in the last six months. Among them, one has type 2 diabetes mellitus and the other does not have any chronic health complications. The unknown etiology controls were SARS-CoV-2 negative, of these, one has hypertension, renal malfunction and died. Three of those four had SARS-CoV-2 antibody-negative tested by All Check COVID-19 IgG/IgM antibody assay kit (CALTH Inc., Republic of Korea) except the deceased patient.

### Metagenomic NGS sequencing

Total nucleic acids were extracted from 300 μL of nasopharyngeal samples and eluted with 60 μL sterile RNase-free water using a commercial kit (Zymo total nucleic acid, USA). Total NA concentration was assayed by Qubit RNA HS Assay Kit (Thermo Fisher Scientific, USA) with Qubit 4 Fluorometer. The extracts were enriched and processed for library preparation using kit Ion Total RNA-Seq Kit v2.0 (Thermo Fisher Scientific, USA) according to the manufacturer’s instructions with a minor modification. In brief, RNA fragmentation was performed using RNase III treatment for 3 minutes at 37°C following the magnetic bead cleanup to optimize the library size at about 200 bp. After adapter ligation of each sample, cDNA was prepared and Ion Xpress™ RNA-Seq Barcode BC primers (Thermo Fisher Scientific, USA) were added. The amplification of the barcoded cDNA was extended to18 cycles instead of 16 due to low concentration nucleic acid in the samples. The final concentrations of the libraries were diluted into 200 picomolar (pM) instead of 100 pM as suggested in the manufacturer’s protocol. One extraction control with sterile water and 11 unknown samples were used for the library preparation. In three mNGS run, four equimolar libraries were pooled for the preparation of template-positive Ion Sphere™ Particles (ISPs) using the Ion 520™ & Ion 530™ Kit – OT2 (Thermo Fisher Scientific, USA) on the Ion One Touch™ 2 System (Thermo Fisher Scientific, USA). Template-positive ISPs were enriched on Ion One Touch™ ES system (Thermo Fisher Scientific, USA). The enriched template-positive ISPs and control ISPs were loaded in Ion 520/530™ chip and sequenced with the next-generation sequencing in the Ion S5™ systems (Thermo Fisher Scientific, USA). The data outputs were analyzed using the automated, streamlined Torrent Suite software (v5.10.0). The primary baseline data were obtained after removing duplicated reads, the average quality scores below Q20, low-quality 3-end reads and adapter sequences.

The CGView Server (http://stothard.afns.ualberta.ca/cgview_server/) was used to construct the circular ring of SARS-CoV-2 genome comparison using the blastn ^13^ and SARS-CoV-2 isolate Wuhan-Hu-1 (NC_045512.2) was used as a reference. Average nucleotide identity (ANI) was calculated using jSpecies ^14^ to compare the SARS-CoV-2 genome with the reference genome.

### Bioinformatics processing and taxonomic assignment

The Binary Alignment Map (bam) files were transferred to FASTQ format through SAMtools ^15^ followed by filtering through BBDuk ^16, 17^ (with options ftm = 5, k = 21, mink = 6, minlen = 30, ktrim = r, qtrim = rl, trimq = 20, overwrite = true) to remove all low-quality sequences. On average, 1.34 million reads per sample (maximum=3.18 million, minimum=0.45 million) passed the quality control step (Supplementary Data 1). The host sequences from the trimmed files were removed by aligning to the human genome (hg38) by using Burrows-Wheeler Aligner (BWA) ^18^ and SAMtools ^15^. The taxonomic assignment has been done by Kraken2 ^19^ with NCBI RefSeq Release 201 database (Bacterial, Viral, Archaeal, and Fungal). Less than 100 hits were not considered for bacteria and fungus, and less than 10 hits were not considered for virus and archea for analysis. Data normalization was performed by previously described methods by multiplying the mean with the proportion ^20^.

### Statistical analyses

The alpha diversity of microbial communities among different groups were compared by calculating the Shannon and Simpson 1-D diversity indexes, Observed, and the Chao-1 richness index using the “Vegan” package in R. The non-parametric test Kruskal-Wallis rank-sum test was used to evaluate alpha diversity and the pairwise Wilcoxon rank-sum test was used to assess pairwise comparison in different groups. Beta-diversity (PCoA) was determined using the Bray– Curtis dissimilarity index, using permutational multivariate analysis of variance (PERMANOVA), to estimate a p-value for differences among the study groups. Phyloseq and vegan packages were employed for those statistical analyses ^21^. Spearman’s correlation coefficient and significance tests were calculated using the R package Hmisc. A correlation network was constructed and visualized with Gephi (ver. 0.9.2). A quantitative analysis of comparative RNA-seq data using shrinkage estimators for dispersion and fold change was employed for differential bacterial species with a statistical significance (q-value) <0.01 and absolute value of log2 (Fold Change) > 3 using DESeq2 (v4.0). The Benjamini-Hochberg correction was used to obtain FDR adjusted p-values (q-values) for multiple hypothesis testing ^22^.

### Determination of protein structures and their binding affinity

The SWISS-MODEL homology modeling webtool and I-TASSER were utilized for generating the three-dimensional (3D) structures of the extracellular toxin protein of the probiotics or outer membrane protein of the pathogen found in our study. We employed CPORT to find out the active and passive protein-protein interface residues of the proteins and peptides. The molecular docking of the bacteriocin and virulent protein of the pathogens were performed using the HADDOCK (v2.4) to evaluate the interaction. The binding affinity of the twenty-four docked complexes was predicted using the PRODIGY.

### Ethical Approval

Ethical approval to conduct this metagenomic study was granted by the Ethical Review Committee of the Jashore University of Science and Technology (ERC no: ERC/FBST/JUST/2020-41). Informed consent was taken from all the COVID-19 positive patients and healthy volunteers.

## Results

All 11 study subjects were divided into three groups, the control group (N=4), SARS-CoV-2 positive group without any history of comorbidity (N=4) and SARS-CoV-2 positive diabetic group. There was no significant difference in age and BMI index in all groups (p-value 0.20 And 0.49 respectively). The detailed patient demography with symptoms, severity, and outcomes were described in detail in supplementary table s1.

### SARS-CoV-2 RNA quantification and genomic data analysis

Metagenomic sequence data analysis retrieved three complete (GISAID Accession ID: EPI_ISL_746318, EPI_ISL_746319 and EPI_ISL_746323) and one partially complete genome sequences of SARS-CoV-2 out of 7 positive samples. Their phylogenetic GISAID clades were GH (Spike D614G, N S194L, NS3 Q57H, NSP2 S263F, NSP3 P74L, NSP12 P323L), G (Spike D614G, N G204R, N R203K, NS3 Q57H, NSP2 I120F, NSP12 P323L, NSP15 R138C) and GR (Spike D614G, E N66B, N G204R, N R203K, NS3 D210B, NS3 P42L, NS6 E55G, NSP2 I120F, NSP3 D1121B, NSP12 N209B, NSP12 P323L, NSP13 L428F). The partial and complete sequences of SARS-CoV-2 were plotted and visualized in a circular ring (Figure s1). The complete genomes were aligned with the reference genome of SARS-CoV-2 isolate Wuhan-Hu-1 (NC_045512.2) and the aligned nucleotide was found 86%, 99% and 98% with the reference genome.

In all seven SARS-CoV-2 positive samples, the average RNA copies were 231,375 among outpatients, 198 among ICU patients and 2,600 among departed patients. No significant relationship was found between the RNA copies and the severity of the disease (Data not shown).

### Microbial diversity and dissimilarity index

The comprehensive assessment of microbial population on the host traits using α-diversity-based association analysis found diverse microbial populations in all three groups of samples, however, they were statistically nonsignificant. The microbial diversity of the control, SARS-CoV-2 non-diabetic and SARS-CoV-2 diabetic groups were non-significant in Shannon (p=0.18), Simpson (p=0.23), Observed (p=0.15) or Chao (p=0.19) (Figure s2). Visualization of community compositions was observed by the Principal Coordinates analysis of Bray-Curtis indicated a significant dissimilarity index in those groups [PERMANOVA: Pseudo-F =1.77, p= 0.047). The control group and SARS-CoV-2 non-diabetic group tended to take a position in the middle of the plot, unlike the SARS-CoV-2 diabetic group. Data from a deceased patient in this co-morbid group was even more scattered. The number of taxonomic units (species) in the control, SARS-CoV-2 non-diabetic and diabetic group were 134, 120, and 162, respectively. More than 14% were shared by all three groups and more than 18% were overlapped between SARS-CoV-2 positive diabetes and non-diabetes group (Figure s3).

### Bacterial diversity and dissimilarity index

The bacterial populations on the host traits were also assessed by α-diversity-based association analysis among the three groups and the Shannon diversity index exhibited significant bacterial diversity (p=0.05). However, Simpson (p=0.09), Observed (p=0.18) or Chao (p=0.18) diversity index differ insignificantly (Figure 1). The Principal Coordinates analysis of Bray-Curtis index PERMANOVA analysis found significant (PERMANOVA: Pseudo-F=2.012, p=0.02) dissimilarities in bacterial species among those groups.

**Figure 1:**
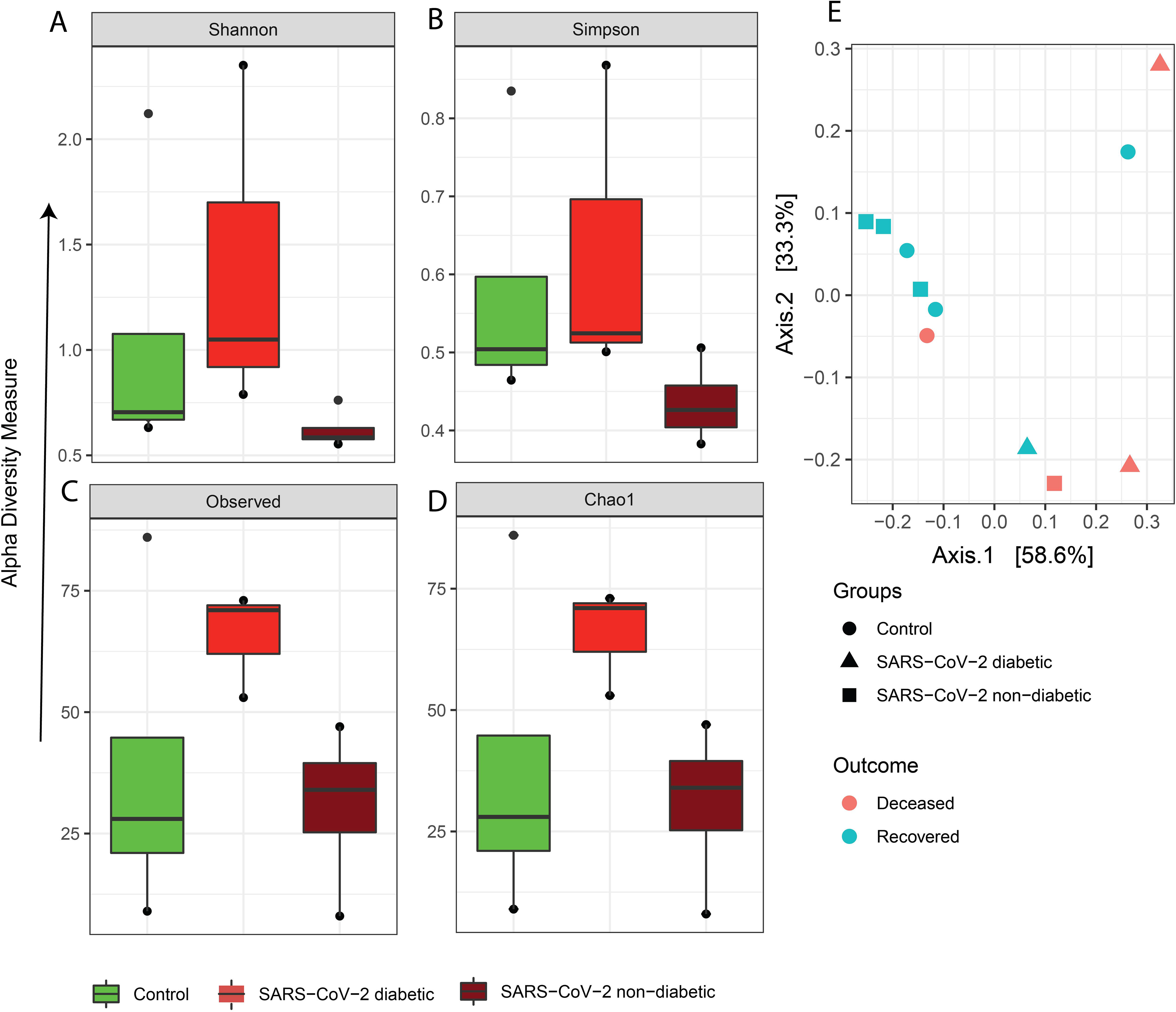
Bacterial α-diversity-based association analysis by (A) Shannon diversity index, (B) Simpson diversity index, (C) observed, and (D) Chao. (E) Principal coordinate analysis by Bray-Curtis dissimilarity index among healthy, recovered and deceased patients.

At phyla level, Firmicutes were the most abundant in all three groups following Bacteroidetes and Proteobacteria. Fusobacteria were abundant only in the control group (Figure s4). This study identified 207 bacterial species among all cases, of which 22 were pathogens, 30 were opportunistic pathogens, 20 were normal-flora, 8 were probiotic and 127 were commensals. The two-way ANOVA analysis found that 41% (9/22) of the pathogens, 47% (14/30) of the opportunistic pathogens, 20% (4/20) of the normal-flora, 25% (2/8) of the probiotics and 20% (25/127) of the commensals were differing significantly between the groups (Table s2).

### Abundance of pathogens, opportunistic pathogens, normal-flora and probiotics

The most abundant species were *Clostridium botulinum, Bacillus cereus, Prevotella melaninogenica, Escherechia coli, Staphylococcus aureus, Prevotella oris, Proteus mirabilis, Pasteurella multocida, Lacrimispora sphenoides, Tennerella forsythia, Salmonella enterica* and *Alkalihalobacillus pseudofirmus* present in all groups. The SARS-CoV-2 positive diabetic group consisted of 41 pathogens/opportunistic pathogens (Figure 2A and 2B). Of which, A*cinetobacter nosocomialis, Shigella flexneri, Bordetella pertussis, Dialister pneumosintes, Sterptococcus orlis, E. fergusonia, Achoromobacter sp. Selenomonas sp., Cutibacterium acnes, Dolosigranulum pigrum, Pseudomonas aeruginosa* and *Stenotrophomonas maltophilia* were present solely in that group. Furthermore, *K. pneumoniae, E. coli* O157:H7, *Yersinia pestis, Porphyromonas* and *Enterobacter* were present in both SARS-CoV-2 positive diabetic and non-diabetic groups. In contrast to that, *Neisseria meningitidis, Haemophillus pittmaniae* and *Streptococcus parasanguinis* were present only in the control group. Moreover, 12 out of 20 species of normal-flora were solely found in the control group, although they were absent in both SARS-CoV-2 positive diabetic and non-diabetic group. Only three species of normal-flora were common in all groups and four species of normal-flora were present in both the control and the SARS-CoV-2 positive diabetic group, the rest was found only in the SARS-CoV-2 diabetic group (Figure 2C). All known probiotic species of *Streptococcus, Lactobacillus, Enterococcus* or *Bifidobacterium* were absent in the control group.

**Figure 2:**
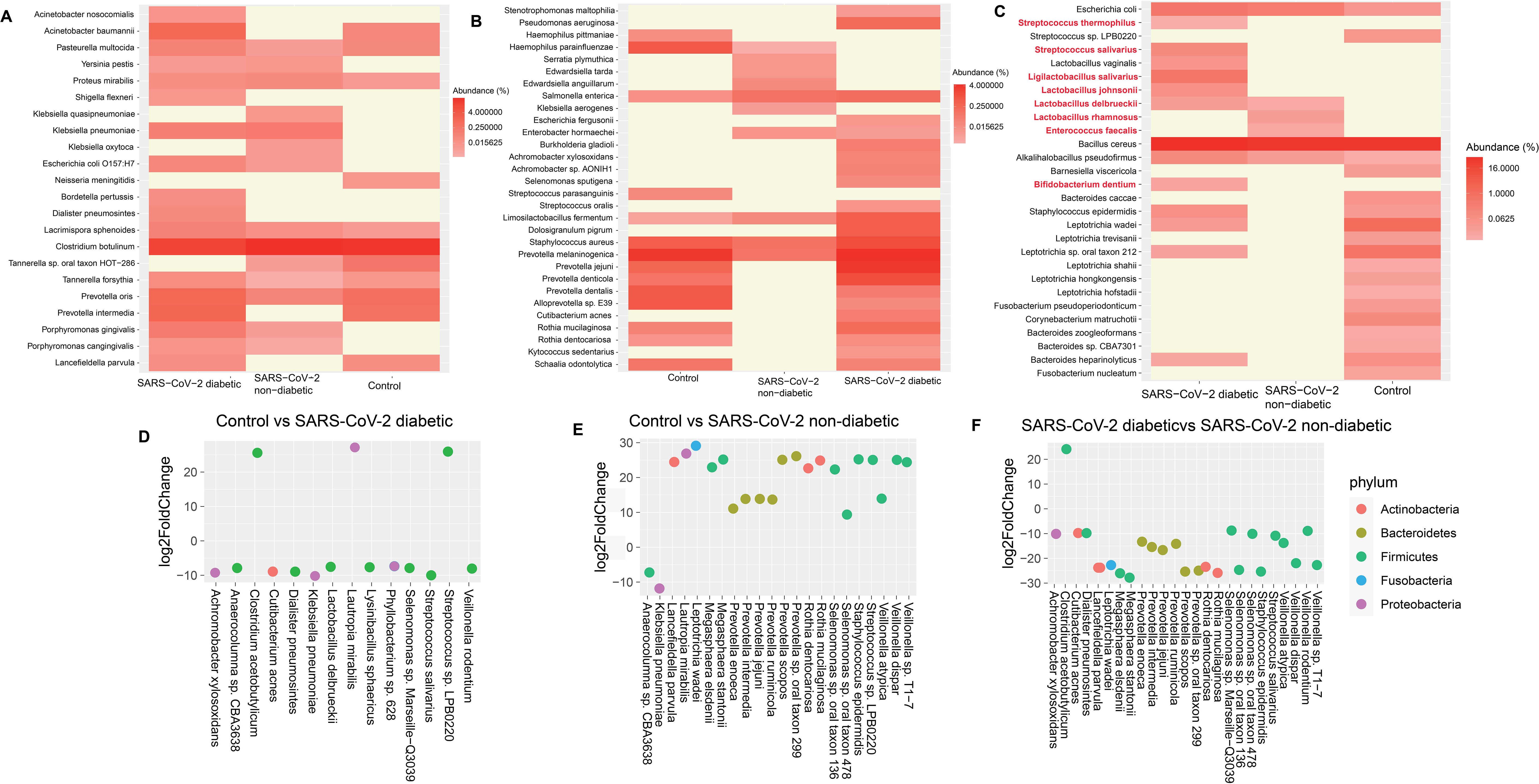
Comparison of bacterial species with relative abundance among the groups of cases. (A) Presence of known pathogenic species with the relative intensity of bacterial genome. (B) Presence of opportunistic pathogen among different groups with relative abundances. (C) The relative abundance of normal-flora and known probiotic species (red highlights). (D) The significant difference of bacterial species and phyla by DESeq2 analysis with log_2_fold changes between control and SARS-CoV-2 diabetic group (p<0.05) (E) The significant difference of bacterial species by DESeq2 analysis with log_2_fold changes between control and SARS-CoV-2 non-diabetic group (p<0.05) (F) The significant difference of bacterial species by DESeq2 analysis with log_2_fold changes between SARS-CoV-2 diabetic group and SARS-CoV-2 non-diabetic group (p<0.05).

The DESeq2 RNA sequence data analysis illustrated the difference of bacterial species between the groups (Figure 2D, 2E and 2F). Several pathogen, opportunistic pathogens and normal-flora differ significantly SARS-CoV-2 diabetic and non-diabetic group compared to control (Benjamini-Hochberg corrected p < 0.05) (Figure 2D, 2E).

### Networking of bacteria

Spearman’s correlation coefficients analyses illustrated 183 significant positive correlations with r range of 0.6 to 1 (P < 0.05) among all pathogen and opportunistic pathogen. *Pseudomonas aeruginosa* positively associated with 14 other pathogenic bacteria including *Dialister pneumosintes, E. coli* O157:H7, *Prevotella intermedia*, *Acinetobacter nosocomialis* and synergistically correlated with *Clostridium botulinum*. *D. pneumosintes* was positively associated with 12 other pathogenic bacteria. The increasing abundance of *Tennerella forsythia* in SARS-CoV-2 diabetic group compared to control was also associated with 11 other pathogenic bacteria. *K. pneumoniae* was positively associated with *Yersinia pestis, E. coli* O157:H7, *Enterobacter sp., S. enterica, Streptococcus oralis*. *H. parainfluenzae*, was positively associated with *H. pittmaniae, N. meningitidis, Alloprevotella sp.* and *Tennerella sp.* A total of 13 significant negative correlations (P < 0.05) were observed associated with *C. botulinum* ranging from −0.67- to −0.77 correlation (r) value.

Correlation analysis (P < 0.05) with probiotics and normal-flora were found to have 109 positive correlations with r > 0.6. Control group featured with a cluster of 15 normal-flora, mostly of *Leptotrichia* and *Bacteroides*, positively associated with each other. In SARS-CoV-2 diabetic group, 6 of normal flora and 6 probiotics were showing significant positive associations (Figure 3B).

**Figure 3:**
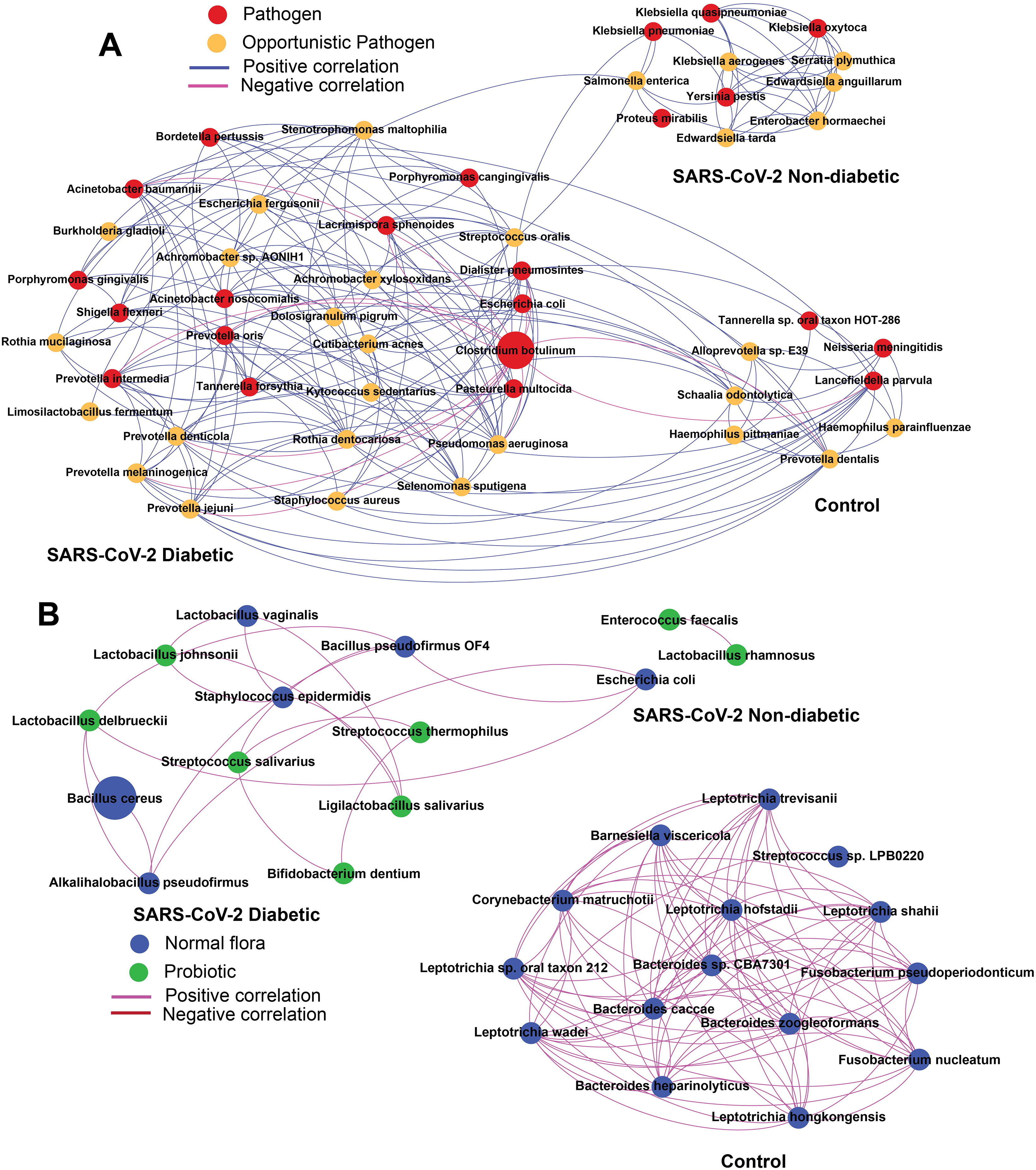
(A) The network analysis showing the co-occurrence patterns of pathogen opportunistic pathogen. Positive spearman correlation represents (r > 0.6) with significant (P < 0.05) correlation and negative Spearman correlation (r) range −0.67- from −0.77with significant (P < 0.05) correlation. The node size is proportional to the mean abundance of the species. (B)The network analysis showing the co-occurrence patterns of probiotics and normal-flora. Positive spearman correlation represents (r > 0.6) with significant (P < 0.05) correlation and no negative correlation was found here. The node size is proportional to the mean abundance of the species.

### Probiotic’s bacteriocins vs pathogen’s outer membrane or toxins

The structure of Salvaricin G32, KLD4 (alpha), Lactacin F, Thermophylin and Lactocin F were generated as released by the specific probiotic bacteria identified in this study (Figure 4). The structure of Enterocin produced by *Enterococcus faecium* and pathogenic proteins were available in the Research Collaboratory for Structural Bioinformatics Protein Data Bank (RCSB PDB, http://rcsb.org) ^23^. All bacteriocins released by those probiotics and pathogenic proteins were showing variable strength of binding affinities (Table 1). The more negative the score the stronger the bond. The KLD4 (alpha) of *Ligilactobacillus salivarius* shows higher binding or neutralizing capacity against Outer membrane protein, Omp33 of *A. baumanii*, Shiga toxin of *E. coli* O157:H7, Hemolysin of the *P. intermedia.* Lactacin F of *Lactobacillus johnsonii* showed the highest binding affinity against the outer membrane protein A of *K. pneumoniae*.

**Table 1:** Binding affinity energy of Bacteriocins from probiotic bacteria with toxins or outer membrane protein of pathogens.

| Bacteriocins | Outer membrane<br>Protein A<br>*Klebsiella pneumoniae* | Outer membrane<br>protein Omp33<br>*Acinetobacter baumannii* | Shiga toxin<br>*E. coli*<br>O157:H7 | Hemolysin<br>*Prevotella*<br>*intermedia* |
| --- | --- | --- | --- | --- |
| Salivaricin G32:<br>*Streptococcus salivarius* | -10.4 | -12.9 | -9.8 | -9.6 |
| KLD4(alpha):<br>*Ligilactobacillus salivarius* | -6.5 | -14.7 | -14.4 | -10.1 |
| LactacinF:<br>*Lactobacillus johnsonii* | -12.1 | -11.1 | -8.4 | -8.1 |
| Thermophylin:<br>*Streptococcus thermophilus* | -9.2 | -12.9 | -9.7 | -6.5 |
| Lactocin 705:<br>*Latilactobacillus* | -8.5 | -12.8 | -10.2 | -8.2 |
| Enterocin:<br>*Enterococcus faecium* | -9.7 | -13.0 | -10.9 | -9.3 |

| Bacteriocin | Outer membrane Protein A<br>*Klebsiella pneumoniae* | Outer membrane protein Omp33<br>*Acinetobacter baumannii* | Shiga toxin<br>*E.coli* O157:H7 | Hemolysin<br>*Prevotella intermedia* |
| --- | --- | --- | --- | --- |
| Salivaricin G32_*Streptococcus salivarius* | -10.4 | -12.9 | -9.8 | -9.6 |
| KLD4 (alpha)_*Ligilactobacillus salivarius* | -6.5 | -14.7 | -14.4 | -10.1 |
| LactacinF_*Lactobacillus johnsonii* | -12.1 | -11.1 | -8.4 | -8.1 |
| Thermophylin *Streptococcus thermophilus* | -9.2 | -12.9 | -9.7 | -6.5 |
| Lactocin 705_ bacteriocin [Lactilactobacillus] | -8.5 | -12.8 | -10.2 | -8.2 |
| Enterocin _*Enterococcus faecium* | -9.7 | -13.0 | -10.9 | -9.3 |

**Figure 4:**
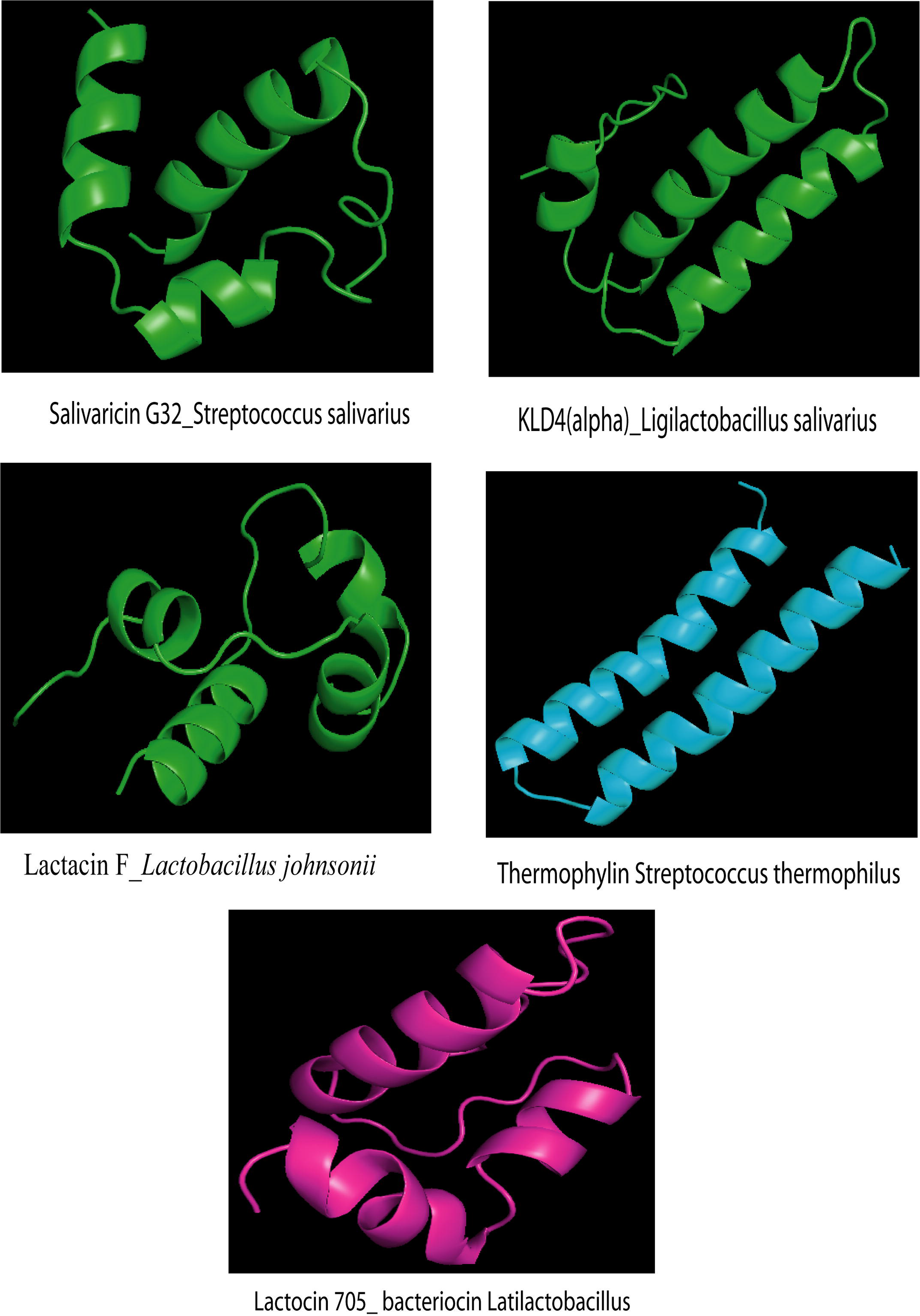
Putative three-dimensional structure of Salvaricin G32, KLD4 (alpha), Lactacin F, Thermophylin and Lactocin F produced by specified probiotic bacteria. The structures were generated using I-TASSER.

## Discussion

SARS-CoV-2 virus infected over 83 million people including 1.8 million deaths globally in 2020. The infections caused by this virus affected mostly people with old age, obesity, type 2 diabetes, hypertension and cardiovascular diseases, especially among males ^24^. Microbiome analysis of co-morbid patients is thus imperative to understanding the influence of microbiota on immune processes in SARS-CoV-2 infections ^25^. Our study compared the baseline nasopharyngeal microbiome of SARS-CoV-2 infected diabetic and non-diabetic patients with controls adjusted with age and gender. The SARS-CoV-2 genome identified among the study enrolled patients belonged to G, GR and GH phylogenetic clades. These three clades cover more than 85% of the strains worldwide ^26^. Our study also analyzed various alpha (α) and beta (β) diversity indexes to observe metagenomic variations in all samples. α-diversity provides the effect of disparity in species to understand microbial communities and diversities on a host. The α-diversity profile for bacteria in our study indicated that the control, SARS-CoV-2 diabetic, and non-diabetic groups have a significant exponential Shannon diversity index (p=0.05). β-diversity provides the index of variation in species composition among different habitats. The β-diversity bray matrix PERMANOVA analysis in this study indicated significant dissimilarities of all microbial communities (p=0.04) and bacterial inhabitants (p=0.02) in all three groups. Firmicutes, Bacteroidetes and Proteobacteria were found abundantly in all three groups. However, Fusobacteria were highly abundant in the control group and Actinobacteria were highly abundant in SARS-CoV-2 diabetic group (Figure s4). The dominance of Firmicutes in diabetic patients were also described in other studies ^27, 28^.

The bacterial species in SARS-CoV-2 diabetic and non-diabetic groups were pathogens enriched compared to control (Figure 2A); however, the SARS-CoV-2 diabetic group was enriched with opportunistic pathogens compared to others (Figure 2B). In this study, the SARS-CoV-2 diabetic group consisted of 18 (82%) pathogens, 23 (77%) opportunistic pathogens, 8 (40%) normal-flora and 6 (75%) probiotics compared to the control group which consisted 12 (55%) pathogens, 14 (47%) opportunistic pathogens, 19 (95%) normal-flora and no (0%) probiotic. SARS-CoV-2 non-diabetic group contained 14 (64%) pathogens, 10 (33%) opportunistic pathogens, 3 (15%) normal-flora and 3 (38%) probiotics. The Kruskal-Wallis significance test of variance demonstrated that the SARS-CoV-2 diabetic patients possess a significantly increased species of pathogens (p<0.05) and opportunistic pathogens (p<0.05) compared to the control and SARS-CoV-2 non-diabetic group. The taxonomic unit (species) of pathogenic bacteria (both pathogens and opportunistic pathogens) in this study were 41 in SARS-CoV-2 diabetic group compared to 26 in the control and 24 in SARS-CoV-2 non-diabetic group (Figure 2A, 2B). A similar finding was reported in a comparative cross-sectional study by ^9^ which demonstrated that the SARS-CoV-2 infected ICU patients harbored more pathogenic bacteria and viruses. Another recent study ^29^ also reported a significantly higher abundance of opportunistic pathogens in SARS-CoV-2 infected patients, such as *Streptococcus*, *Rothia*, *Veillonella*, and *Actinomyces*; and a lower relative abundance of beneficial symbionts compared with healthy human. In our study, the controls were enriched with numerous species of normal-flora compared to both SARS-CoV-2 positive groups (Figure 2C). The control group contained 19 species of normal-flora which was reduced to 8 in SARS-CoV-2 diabetic group and only 3 in SARS-CoV-2 non-diabetic group. A group of researchers reported that the specific intestinal microbiota of COVID-19 patients ^30^ could suppress the SARS-CoV-2 attachment. The severe patients might have featured dysbiosis and the normal microbiota has been replaced with pathogenic bacteria ^31^. Hyperglycemia, inflammation and severe oxidative stress in a patient’s physique may alter the oral microbiome ^32^. Other evidence also suggested an association of dysbiosis of the normal microbiota due to diabetes ^33, 34^. Moreover, studies on SARS viral epidemic demonstrated that co-infection was one of the major complications in prolonged hospitalization and mechanical ventilation. Pathogenic bacteria like *E. faecalis, K. pneumonia, A. baumannii*, and *Stenotrophomonas maltophilia* inhabited inside the oral cavity can also cause nosocomial infections ^35^. Several studies found that opportunistic pathogens were the most common cause of secondary infections in viral epidemic ^36, 37^. Pathogenic bacteria like *Legionella pneumophila* ^38^, *N. meningitidis, Moraxella catarrhalis* ^39^ were known to be associated with influenza co-infection. *Porphyromonas gingivalis*, also found in our study, was an important cause of periodontitis ^40^. The use of antibiotic to prevent secondary infections may also lead to the loss of normal-flora and probiotics causing dysbiosis in patients with COVID-19.

One interesting finding in our study was the absence of *H. pneumoniae, N. meningitidis, S. parasanguinis* and *H. pittmaniae* in the SARS-CoV-2 diabetic groups, unlike other pathogenic bacteria (Figure 2A). However, those bacteria significantly enriched the control groups in our study which was also evident by Qin et al ^34^ and his research team. Another interesting feature in our study was the significant reduction of normal-flora in both SARS-CoV-2 patient groups such as *Leptotrichia, Bacteroides, Fusobacterium, Chorynebacterium* and *Bernesiella* spp., indicating the imbalance of microbiota (Figure 2D-2F). Moore et al ^41^ also found a significant reduction of *Fusobacterium periodonticum* in the nasopharynx during SARS-CoV-2 infections. Those bacterial flora found in the oral cavity may inhibit pathogenic bacteria by producing antimicrobial substances such as bacteriocins, lactic acid and hydrogen peroxide which might create a hostile condition for the pathogenic bacteria ^30^. The presence of probiotic species with several pathogens and opportunistic pathogens in the SARS-CoV-2 diabetic group revealed that those probiotics might assist the host by inhibiting those pathogenic bacteria. Our docking analysis provided an evidence of this hypothesis. The binding affinities indicate the bacteriocins from these probiotic bacteria had a strong binding affinity with the pathogenic toxins or outer membrane proteins suggesting that they have the capacity to inhibit the pathogens (Table 1).

The correlation coefficient network analysis in this study found significant positive associations (N=183) and few negative associations (N=13) among the pathogenic bacteria in SARS-CoV-2 infected diabetic patients. The notable associations were observed in SARS-CoV-2 diabetic group with 32 species of pathogenic bacteria (both pathogens and opportunistic pathogens) compared to 11 species in SARS-CoV-2 non-diabetic and 8 species in control group (Figure 3A). This analysis indicated various patterns of pathogenic networks in theSARS-CoV-2 diabetic group especially among enteric pathogens, nosocomial bacteria and other opportunistic pathogens. However, this analysis found no correlation in the most abundant species between the control group and SARS-CoV-2 positive non-diabetic patients.

The co-occurrences of network among the probiotic and normal-flora identified several significant positive associations (N= 109) but no synergistic correlations. There was a separate cluster of 15 normal-flora in the control group with 90 significant positive associations which were absent in SARS-CoV-2 positive diabetic and non-diabetic groups (Figure 3B). The decrease of normal-flora in the later groups indicated that they were outnumbered by the pathogenic species mentioned above. The increase of pathogenic environment results in a non-productive busy immune response in the host which ultimately suppresses the adaptive immune response against SARS-CoV-2. Nevertheless, increased species of probiotic strains (N=6) in the SARS-CoV-2 diabetic group compared to the control group (N=0) and SARS-CoV-2 non-diabetic group (N=3) indicated an inadequate resistance against highly abundant pathogenic bacteria. Another study ^42^ demonstrated that immunomodulatory probiotics, *Rothia mucilaginosa, K. oxytoca, Enterobacter kobei, Bacillus cereus, Faecalibacterium prausnitzii* etc. were enriched at COVID-19 positive patients. Our study had a limitation that the microbiome analyses was performed with small number of individuals studied. In developing countries, this limitation is quite common because of paradoxical situations; doubled price of metagenome reagents in developing countries, extended delivery time with short period of expiry, and unavailability of reagents during the peak times of COVID-19 infections. Therefore, there are very few data reported from those regions where the highest number of patients are having comorbidity. However, our preliminary observations and hypothesis were supported by appropriate statistical methods and the results are compared with suitable controls.

## Conclusions

The SARS-CoV-2 positive diabetic patients were possessed by significantly increased pathogenic species compared to the control and SARS-CoV-2 non-diabetic group. In both groups, the normal-flora strains were replaced by numerous pathogenic bacterial species which might correlate the severity and outcome of complications. Patients within the SARS-CoV-2 positive non-diabetic group exhibited significantly increased opportunistic pathogens compared to the control. Those dysbiosis suppressed the adaptive immune response against SARS-CoV-2 because of induced immune response against those pathogenic bacteria. Presence of few probiotic species among the SARS-CoV-2 diabetic patients indicated that those probiotics were inhibiting the pathogens as observed in our study. However, the numbers might not be competitive enough to provide successful protection as seen within deceased patients. One approach for maintaining a healthy microbiome in SARS-CoV-2 diabetic patients might include promoting probiotics and normal-flora by dietary changes and reducing pro-inflammatory states. Relocation of the microbial balance with normal-flora and sufficient probiotics may prevent pathogenic environments and excessive inflammations that might enhance the adaptive immune response, leading to better outcomes for the SARS-CoV-2 diabetic patients.

