## supplementary for "Metagenomic diagnosis and pathogenic network profile of SARS-CoV-2 in patients co-morbidly affected by type 2 diabetes"

Table s1: Demographics and clinical characteristics of the study subjects.

| ID | Cases | Groups | Age/ Sex | Sign and Symptoms appeared | SARS-CoV-2 tests results | Chronic health complications |
|---|---|---|---|---|---|---|
| MHC10 | Healthy | Control | 57/ M | No | Antibody negative, rt-PCR negative | None |
| MHC15 | Healthy | Control | 75/ M | No | Antibody negative, rt-PCR negative | Diabetes |
| MOE14 | Other unknown etiology | Control | 57/ M | Feverish (99F) | Antibody negative, rt-PCR negative | None |
| MDB11 | Death | Control | 79/ M | Fever, Dry cough, loss of appetite, pain or swelling of legs, hands | rt-PCR negative | Hypertension, renal impairment |
| MMS2 | Outpatients | SARS-CoV-2 non-diabetic | 33/ M | Dry cough | rt-PCR positive | None |
| MICU1 | ICU | SARS-CoV-2 non-diabetic | 35/ M | Fever; headache; Shortness of breath; difficulty breathing; loss of taste; loss of smell. | rt-PCR positive | None |
| MICU13 | ICU, coma | SARS-CoV-2 non-diabetic | 75/ M | Lung co-infection, Sore throat, difficulty breathing, loss of appetite, loss of taste, loss of smell | rt-PCR positive | None |
| MDN6 | Death | SARS-CoV-2 non-diabetic | 59/ M | Fever, Dry cough, Shortness of breath, pain or swelling of legs, hands | rt-PCR positive | None |
| MHD7 | Hospitalized | SARS-CoV-2 diabetic | 56/ M | Fever, headache, Running nose, Dry cough, Sore throat, Muscle pain, Shortness of breath, loss of taste, loss of smell, Diarrhea | rt-PCR positive | Diabetes, Hypertension |
| MDD8 | Death | SARS-CoV-2 diabetic | 60/ M | Fever, Sore throat, Muscle pain, Shortness of breath, difficulty breathing, loss of taste, loss of smell | rt-PCR positive | Diabetes |
| MDD9 | Death | SARS-CoV-2 diabetic | 65/ M | Fever, bedsore, broken hip | rt-PCR positive | Diabetes |

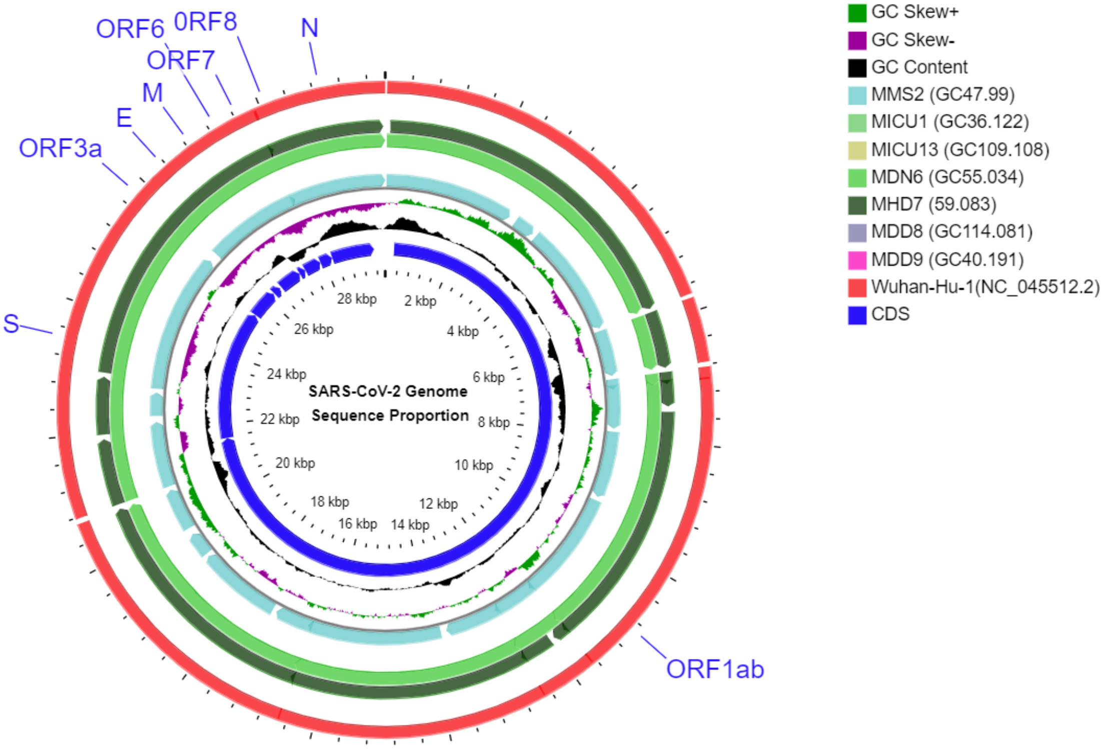


Figure s1: Comparison of the SARS-CoV-2 reference genome Wuhan-Hu-1 (NC_045512.2) with the reads retrieved from metagenomic data using blastn. The CGView Server (http://stothard.afns.ualberta.ca/cgview_server/) was used to construct the ring.

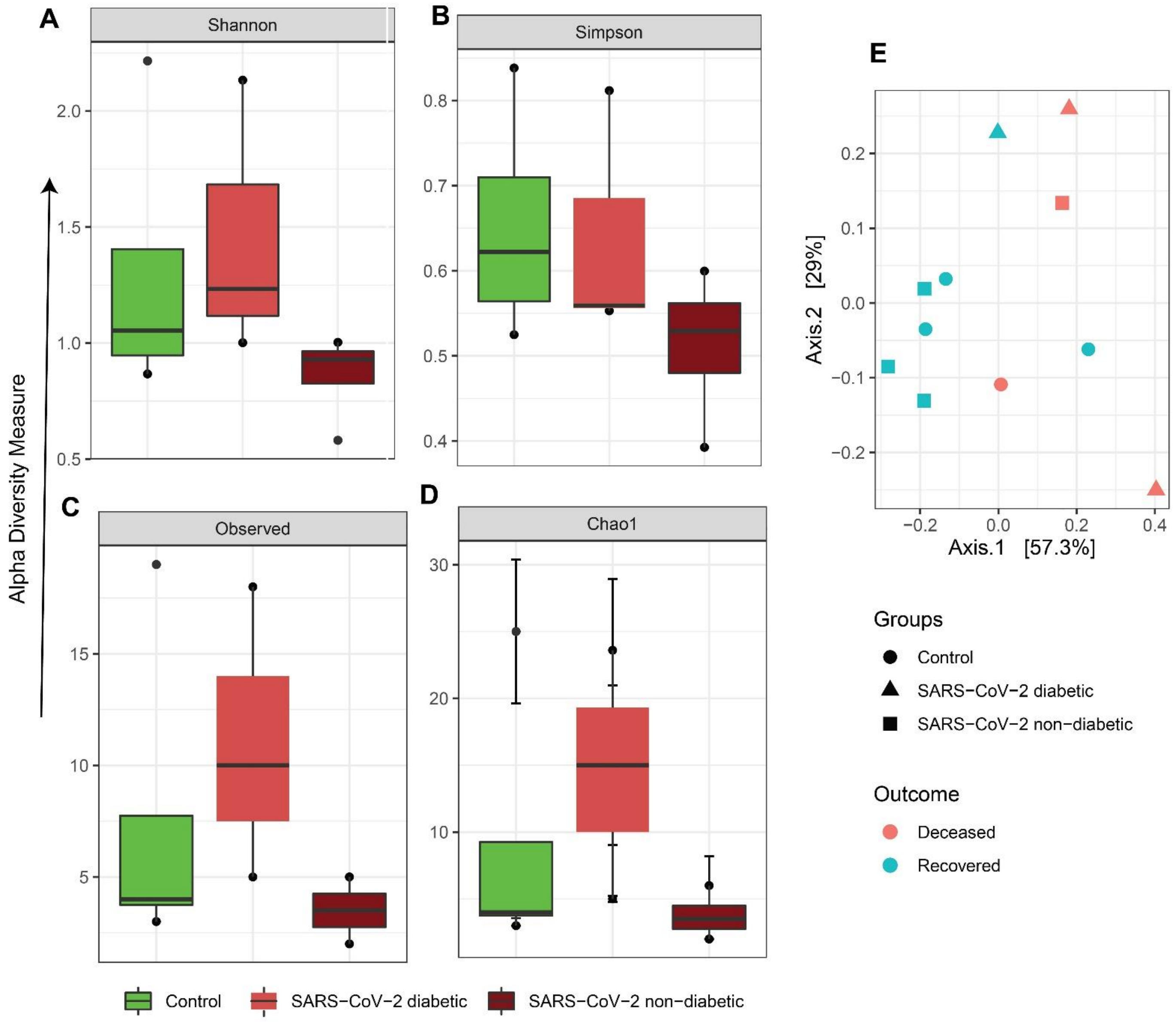


Figure s2: Microbial α-diversity-based association analysis by (A) Shannon diversity index, (B) Simpson diversity index, (C) observed, and (D) Chao. (E) Principal coordinate analysis by Bray-Curtis dissimilarity index among healthy, recovered and deceased patients.

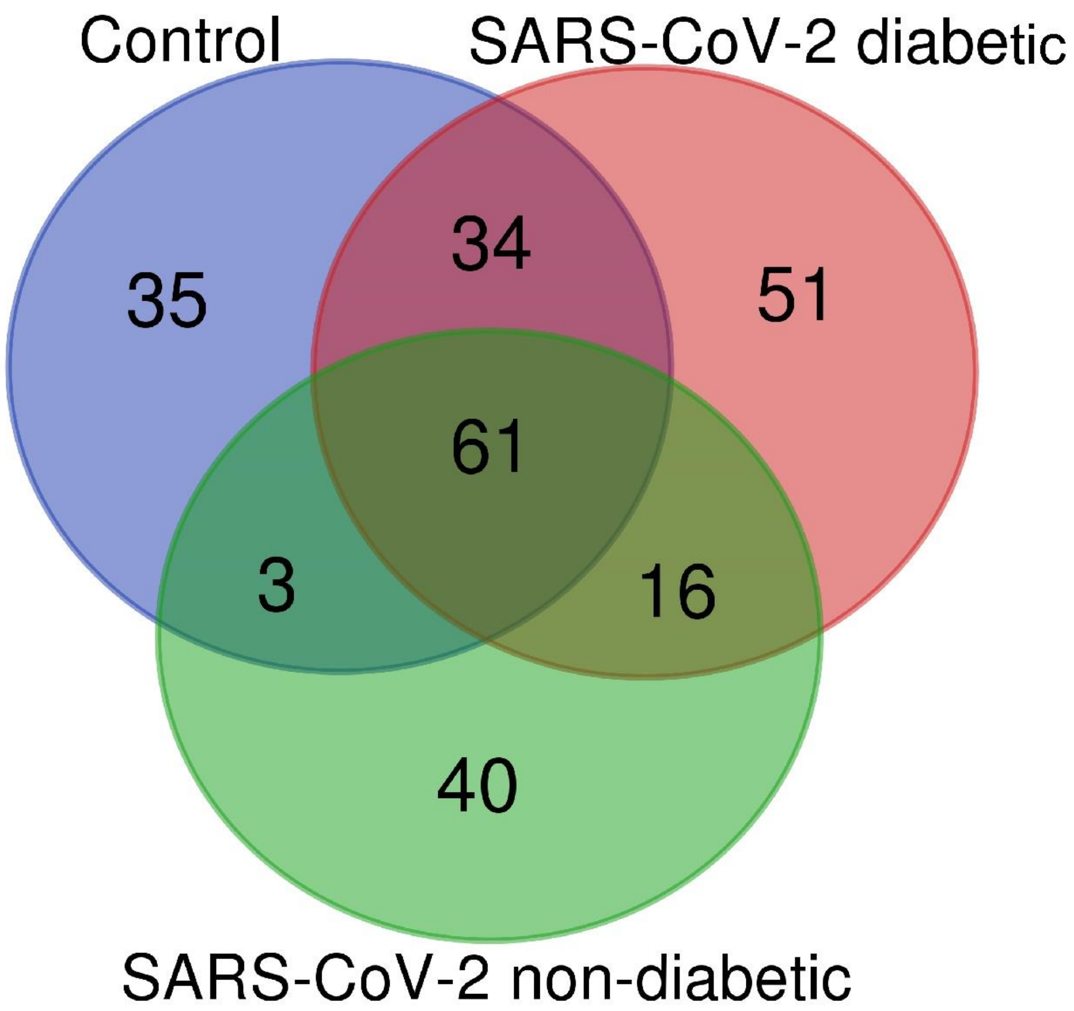


Figure s3: Veen diagram of microbial taxonomic unit (species) among the control, SARS-CoV-2 diabetic and SARS-CoV-2 non-diabetic group.

Table s2: Two-way analysis of variance significance test performed between two groups among the control, SARS-CoV-2 positive diabetic and non-diabetic group.

| Name of the species | Two-way ANOVA significance test (p-value) | | |
|---|---|---|---|
| | Control vs. SARS-CoV-2 non-diabetic | Control vs. SARS-CoV-2 diabetic | SARS-CoV-2 non-diabetic vs. SARS-CoV-2 diabetic |
| *Acinetobacter baumannii* | 0.040 | 0.293 | 0.003 |
| *Atopobium parvulum* DSM 20469 | 0.056 | 0.676 | 0.029 |
| *Clostridium sphenoeides* JCM 1415 | 0.205 | 0.003 | 0.066 |
| *Dialister pneumosintes* | 1.000 | 0.018 | 0.018 |
| *Escherichia coli* O157:H7 | 0.371 | 0.001 | 0.008 |
| *Klebsiella pneumoniae* | 0.001 | 0.000 | 0.456 |
| *Porphyromonas gingivalis* | 0.293 | 0.092 | 0.478 |
| *Prevotella intermedia* | 0.000 | 0.090 | 0.000 |
| *Prevotella oris* | 0.011 | 0.157 | 0.000 |
| *Proteus mirabilis* | 0.049 | 0.137 | 0.733 |
| *Achromobacter xylosoxidans* | 1.000 | 0.023 | 0.023 |
| *Alloprevotella sp.* E39 | 0.013 | 0.304 | 0.204 |
| *Cutibacterium acnes* | 1.000 | 0.012 | 0.012 |
| *Dolosigranulum pigrum* | 1.000 | 0.067 | 0.067 |
| *Haemophilus parainfluenzae* | 0.105 | 0.022 | 0.425 |
| *Limosilactobacillus fermentum* | 0.044 | 0.001 | 0.145 |
| *Prevotella dentalis* | 0.017 | 0.768 | 0.013 |
| *Prevotella denticola* | 0.002 | 0.071 | 0.000 |
| *Prevotella jejuni* | 0.000 | 0.036 | 0.000 |
| *Prevotella melaninogenica* | 0.035 | 0.072 | 0.000 |
| *Pseudomonas aeruginosa* | 1.000 | 0.000 | 0.000 |
| *Rothia dentocariosa* | 0.389 | 0.196 | 0.037 |
| *Rothia mucilaginosa* | 0.056 | 0.960 | 0.085 |
| *Salmonella enterica* | 0.001 | 0.001 | 0.807 |
| *Schaalia odontolytica* | 0.214 | 0.229 | 0.019 |
| *Staphylococcus aureus* | 0.760 | 0.002 | 0.005 |
| *Lactobacillus delbrueckii* | 0.083 | 0.027 | 0.546 |
| *Ligilactobacillus salivarius* | 1.000 | 0.063 | 0.063 |
| *Streptococcus salivarius* | 1.000 | 0.008 | 0.008 |
| *Bacillus pseudofirmus* OF4 | 0.188 | 0.001 | 0.047 |
| *Escherichia coli* | 0.000 | 0.000 | 0.762 |
| *Streptococcus sp.* LPB0220 | 0.061 | 0.083 | 1.000 |
| *Leptotrichia wadei* | 0.006 | 0.218 | 0.194 |
| *Staphylococcus epidermidis* | 0.293 | 0.144 | 0.015 |
| *Acinetobacter bereziniae* | 0.192 | 0.333 | 0.030 |
| *Alkalihalobacillus pseudofirmus* | 0.188 | 0.001 | 0.047 |
| *Anaerocolumna sp.* CBA3638 | 0.061 | 0.021 | 0.560 |
| *Campylobacter concisus* | 0.023 | 0.035 | 1.000 |
| *Clostridium acetobutylicum* | 0.081 | 0.011 | 0.345 |
| *Corynebacterium segmentosum* | 0.959 | 0.001 | 0.001 |
| *Halomonas sp.* JS92-SW72 | 0.745 | 0.092 | 0.047 |
| *Lacrimispora sphenoides* | 0.205 | 0.003 | 0.066 |

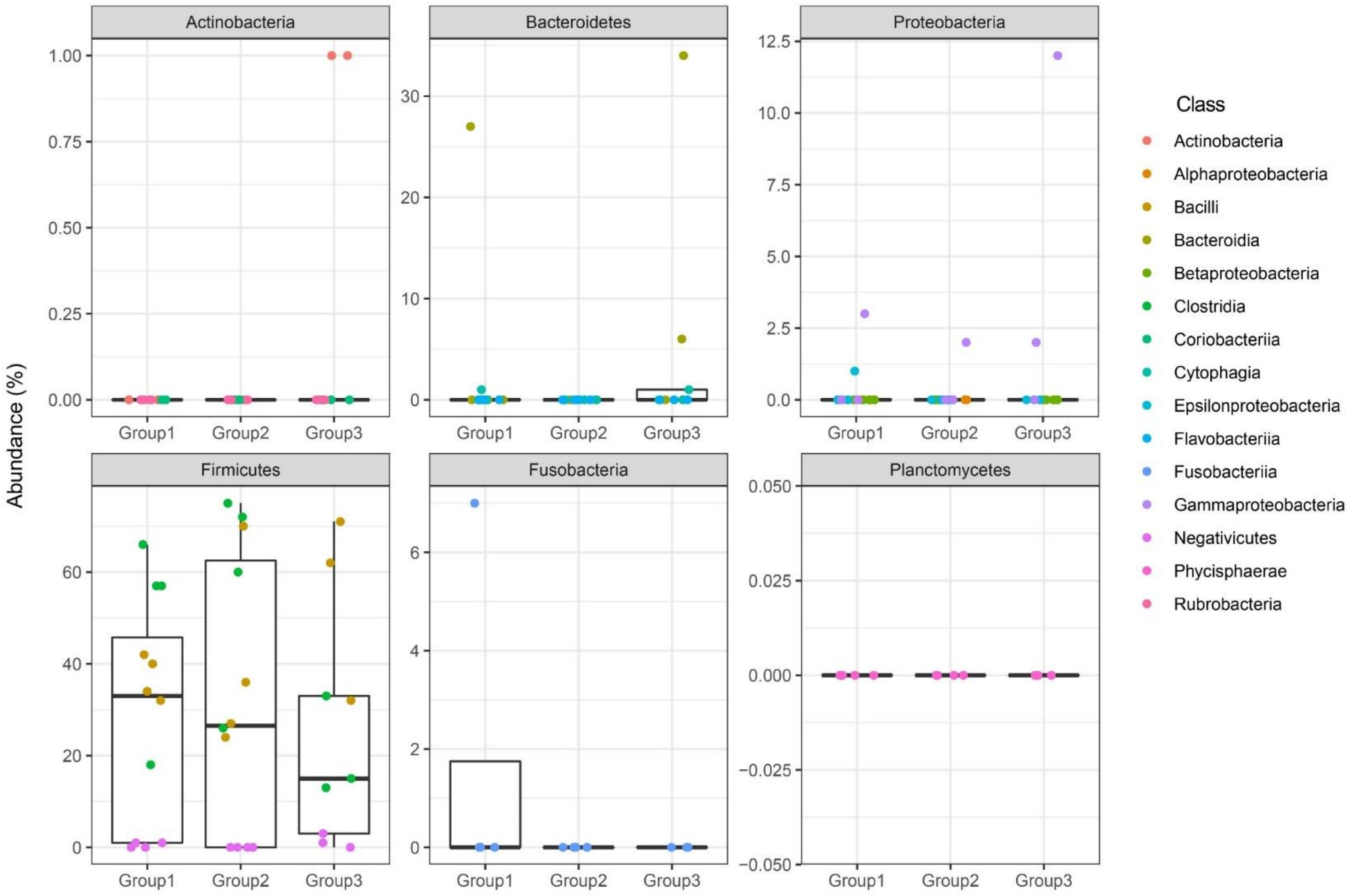


Figure s4: Relative abundance of bacterial taxa within 6 phyla among the control (Group 1), SARS-CoV-2 positive diabetic (Group 2) and SARS-CoV-2 positive non-diabetic (Group 3) group. Data were normalized by the percentage of the total taxonomic unit identified.
